# Effect of fluoxetine on adult amblyopia: a placebo-controlled study combining neuroplasticity-enhancing pharmacological intervention and perceptual training

**DOI:** 10.1101/327650

**Authors:** Henri J. Huttunen, J. Matias Palva, Laura Lindberg, Satu Palva, Ville Saarela, Elina Karvonen, Marja-Leena Latvala, Johanna Liinamaa, Sigrid Booms, Eero Castrén, Hannu Uusitalo

## Abstract

Amblyopia is a common visual disorder that is treatable in childhood. However, therapies have limited efficacy in adult patients with amblyopia. Fluoxetine can reinstate early-life critical period-like neuronal plasticity and has been used to recover functional vision in adult rats with amblyopia. This phase 2, randomized, double-blind (fluoxetine vs. placebo), multicenter clinical trial examined whether or not fluoxetine can improve visual acuity in amblyopic adults. This interventional trial included 42 participants diagnosed with moderate to severe amblyopia. Subjects were randomized to receive either 20 mg fluoxetine (n=22) or placebo (n=20). During the 10-week treatment period, all subjects performed daily computerized perceptual training and eye patching. There was no significant difference in treatment efficacy between the groups. Visual acuity at the primary endpoint had significantly improved over baseline in both the fluoxetine (−0.167 logMAR) and placebo (−0.194 logMAR) groups (both p < 0.001). Because patching alone is not effective in adults, the visual acuity improvement likely resulted from perceptual training. There was a positive correlation between visual acuity improvement and the perceptual training time. While this study failed to provide evidence that fluoxetine enhances neuroplasticity, our data support the usefulness of perceptual training for vision improvement in adults with amblyopia.

## INTRODUCTION

Amblyopia is a condition in which the best-corrected visual acuity (BCVA) is impaired in one eye or, less frequently, both eyes, even though no ocular abnormalities are generally present. Amblyopia develops when one or both eyes have abnormal visual input (either physical or physiological) during the sensitive period in childhood (from birth to 6 years).^1, 2^ Amblyopia is most commonly caused by strabismus in very young children (<3 years of age, 82%)^3^ and by refractive errors (anisometropia, isoametropia) in older children. ^4, 5^ In the clinical setting, moderate amblyopia is defined as a logarithm of the minimum angle of resolution (logMAR) BCVA of ≥0.30 (Snellen equivalent: ≤20/40) and/or an interocular BCVA difference of 0.2 logMAR or more (in cases of unilateral amblyopia). The prevalence of amblyopia in the general population varies from 1.3% to 3.6%, and it is one of the most common causes of monocular visual impairment in adults.^4, 6, 7^ Patients with monocular amblyopia have a significantly increased risk of visual impairment if vision in their “good” eye is lost as a result of trauma or disease.^8, 9^

Amblyopia can be treated in early life, ^10–12^ but visual gains diminish in school-aged children because of a decline in visual system neuroplasticity and, possibly, treatment compliance.^1, 13–15^ Amblyopia is very difficult, if not impossible, to treat in adults.^16^

An improved understanding of the neuronal mechanisms underlying amblyopia and adult brain neuroplasticity^17–19^ has led to the development of visual rehabilitation methods that can be used after the critical period.^20–24^ Experimental models of amblyopia are based on the effects of monocular deprivation on the structure and function of the visual cortex.^25, 26^ Using an animal model, Espinosa and Stryker^26^ showed that the effects of amblyopia can be reversed during the critical period in early postnatal development, but not later in life. However, recent evidence indicates that environmental enrichment^27, 28^ and pharmacological treatment^29^ can reactivate critical period-like plasticity in the visual cortex of adult rodents. In particular, fluoxetine, a selective serotonin reuptake inhibitor (SSRI), promotes neuroplasticity and neurogenesis^30^ and reactivates critical period-like plasticity in the rat visual cortex.^18^

An emerging body of literature suggests that vision improvements can be achieved with conventional therapies (e.g., occlusion therapy) in adolescents (i.e., older children and teenagers)^31–33^ and adults^20, 21, 34, 35^ with amblyopia even though post-childhood amblyopia is generally considered untreatable.

In addition, catecholamine-based medical treatments can temporarily improve vision in human amblyopic patients, including adults.^1, 36, 37^ Perceptual training^38^, dichoptic non-action and action videogame use^39^, and videogame use during patching^40^ can improve vision in the amblyopic eye and binocular vision in adults. This placebo-controlled study examined whether fluoxetine can enhance neuroplasticity and improve vision in adults with amblyopia. The treatment period included eye patching and computerized perceptual training on a web-based system for all subjects.

## RESULTS

### Study subjects

A total of 42 subjects were enrolled in the study, with 22 and 20 subjects randomly assigned to the fluoxetine and control group, respectively. Table 1 presents a complete list of the eligibility and exclusion criteria used for patient selection. Forty-one of 42 subjects (97.6%) required new spectacles before randomization. Four subjects were non-compliant, 3 subjects withdrew their consent, and 1 subject was lost to follow-up. Therefore, a total of 37 subjects ultimately completed the 10-week treatment period, including primary endpoint assessments, and 34 completed the 3-month post-treatment follow-up period (20 in the fluoxetine group and 14 in the control group; Figure 1). Data from all 42 randomized subjects were subjected to an intention-to-treat analysis and were included in analyses. Subjects that completed the study showed good medication compliance (>85%) and completed >85% of the computerized perceptual training sessions.

**Table 1.**
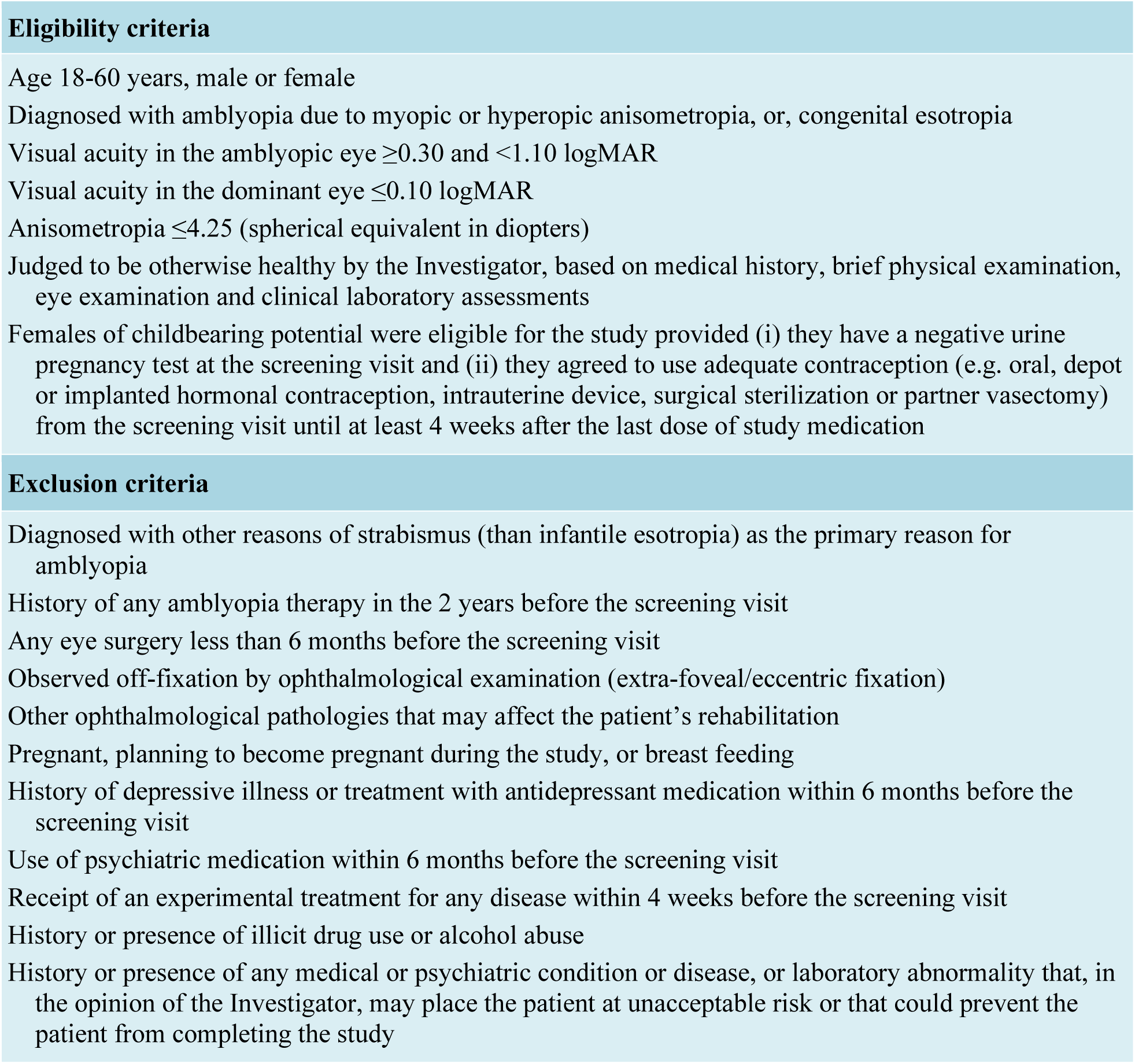
Eligibility and exclusion criteria.

**Figure 1.**
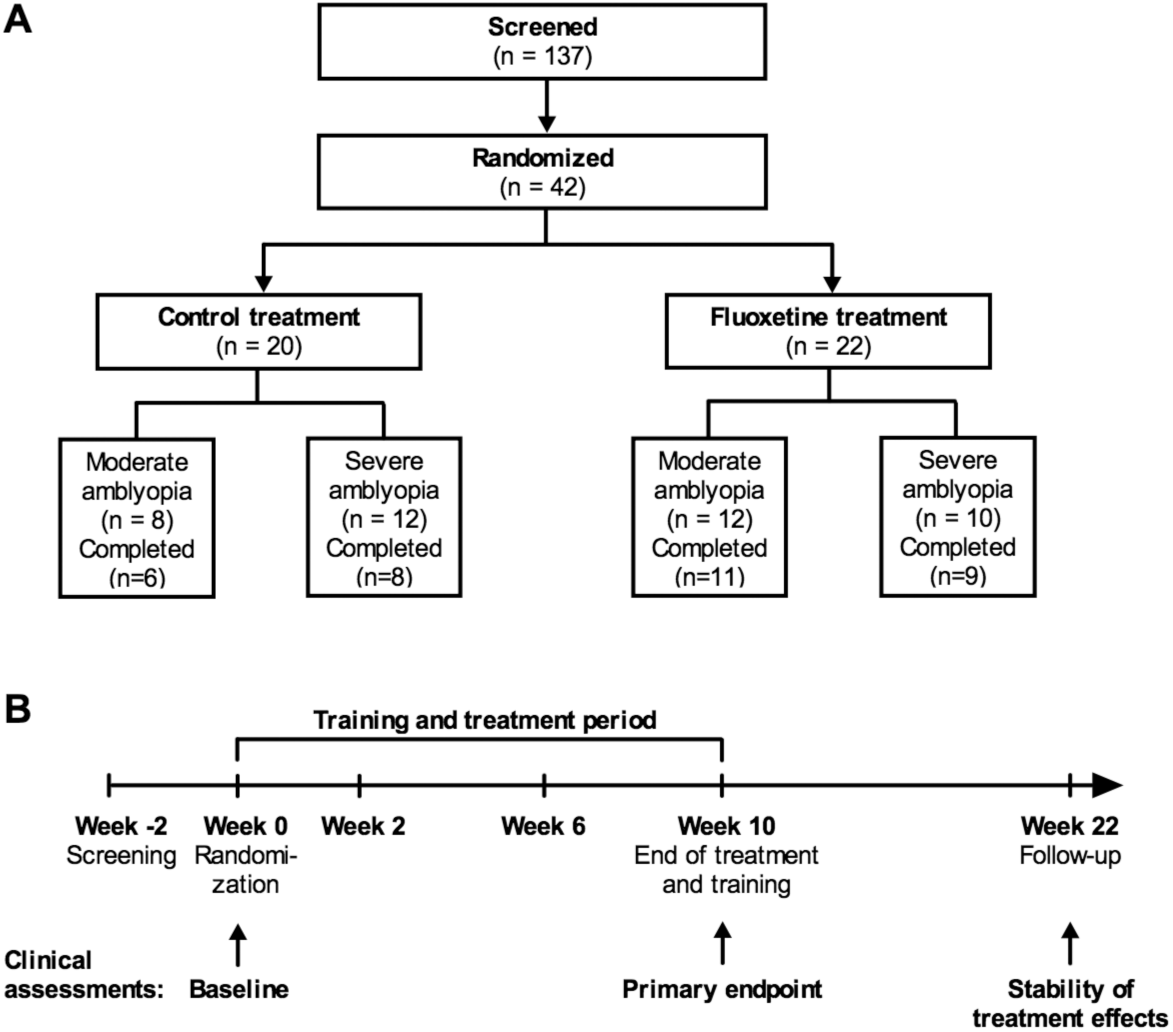
Study design and flow. (A) Disposition and patients. (B) Visit and assessment schedule and duration of medication and active training.

Baseline demographic and ocular parameters are summarized in Table 2. Briefly, best-corrected logMAR visual acuity at baseline was between 0.30 and 1.08 in the amblyopic eye and not worse than 0.10 in the dominant eye. The majority of subjects had anisometropic amblyopia [19 of 20 control subjects (95.0%), 18 of 22 fluoxetine subjects (81.8%)]. Four subjects (1 in the control group and 3 in the fluoxetine group) exhibited combined strabismic-anisometropic amblyopia, while 1 subject in fluoxetine group exhibited strabismic amblyopia. These 5 subjects had undergone strabismus surgery during childhood. Twenty subjects (8 in the control group and 12 in the fluoxetine group) had moderate amblyopia and 22 (12 in the control group and 10 in the fluoxetine group) had severe amblyopia (Figure 1A).

**Table 2.**
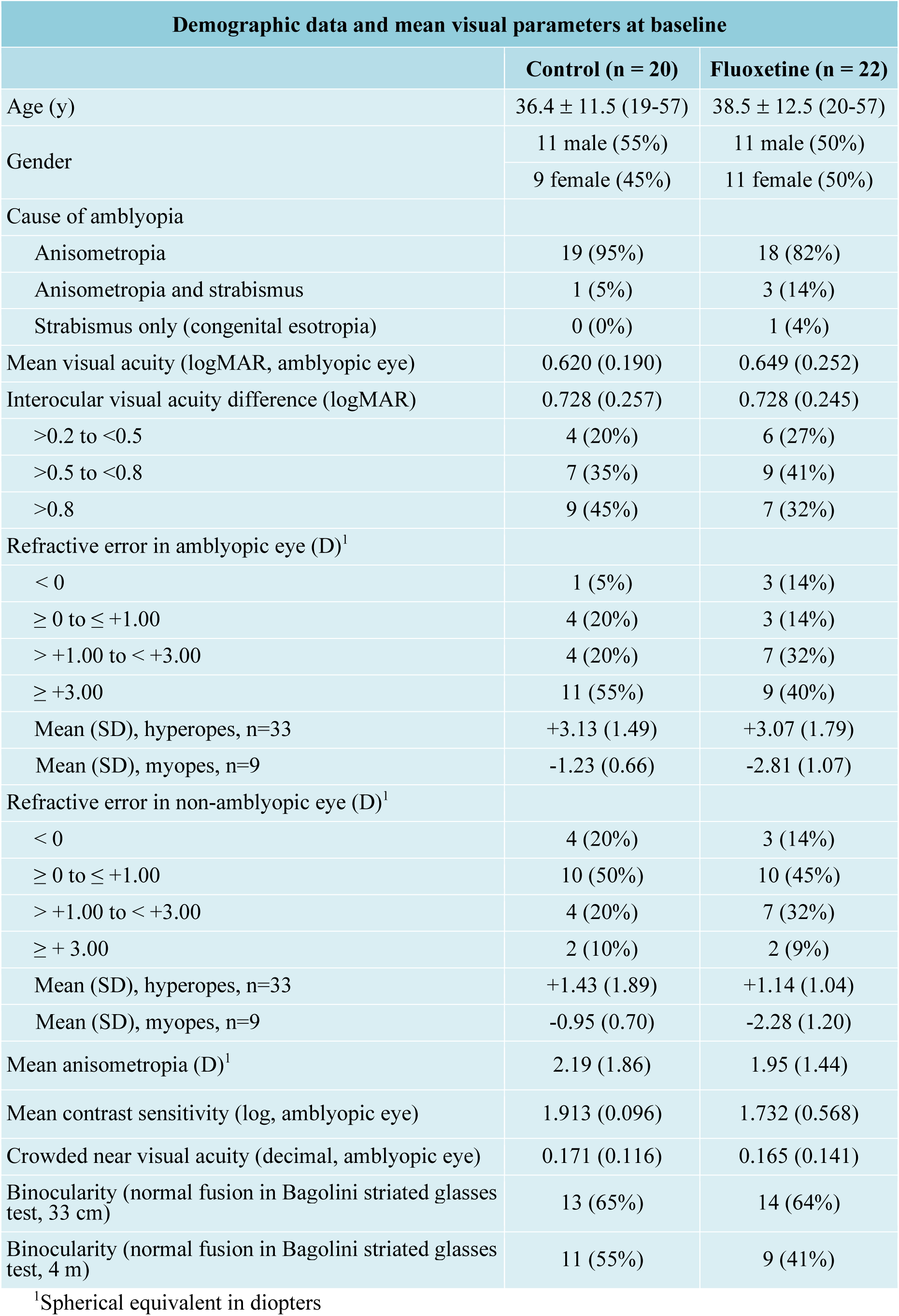
Demographic data and mean visual parameters at baseline.

Mean subject age was 36.4 ± 11.5 years in the control group and 38.5 ± 12.5 years in the fluoxetine group. Mean baseline logMAR visual acuity in the amblyopic eye was 0.620 ± 0.190 (Snellen equivalent: 20/83) in the control group and 0.649 ± 0.252 (20/89) in the fluoxetine group. Overall, the average difference in the logMAR visual acuity between eyes was 0.728 in both groups. For hyperopic subjects (n=33), the mean refractive error in the amblyopic eye was +3.13 ± 1.49 and +3.07 ± 1.79 D in the control and fluoxetine groups, respectively, and for myopic subjects (n=9), the mean refractive error in the amblyopic eye was −1.23 ± 0.66 and −2.81 ± 1.07 D, respectively. Mean anisometropia was 2.19 ± 1.86 and 1.95 ± 1.44 D in the control and fluoxetine groups, respectively. Baseline binocular vision testing revealed that 9 of 20 control group subjects (45.0%) and 13 of 22 fluoxetine group subjects (59.1%) had suppression or anomalous retinal correspondence (ARC) at the 4-m distance. At the 33-cm distance, 7 of 20 subjects in the control group (35.0%) and 8 of 22 subjects in the fluoxetine group (36.4%) showed suppression or ARC at baseline. All subjects had impaired near visual acuity (with crowding effect; near logMAR visual acuity < 0.7) at baseline, but only 2 control group subjects (10.0%) and 6 fluoxetine group subjects (27.3%) had abnormal contrast sensitivity at the intermediate spatial frequency measured using the Pelli-Robson chart. Mean baseline visual parameters are summarized in Table 2.

The 10-week treatment regimen included a combination of medication, eye patching, and perceptual training. A game-based perceptual training software was specifically developed for enhancing the use of the amblyopic eye during patching. The game tasks are illustrated in Figure 2 and Supplementary video 1. The study was designed to include a placebo control group for the medication only, and all participating subjects followed the same daily patching and training instructions.

**Figure 2.**
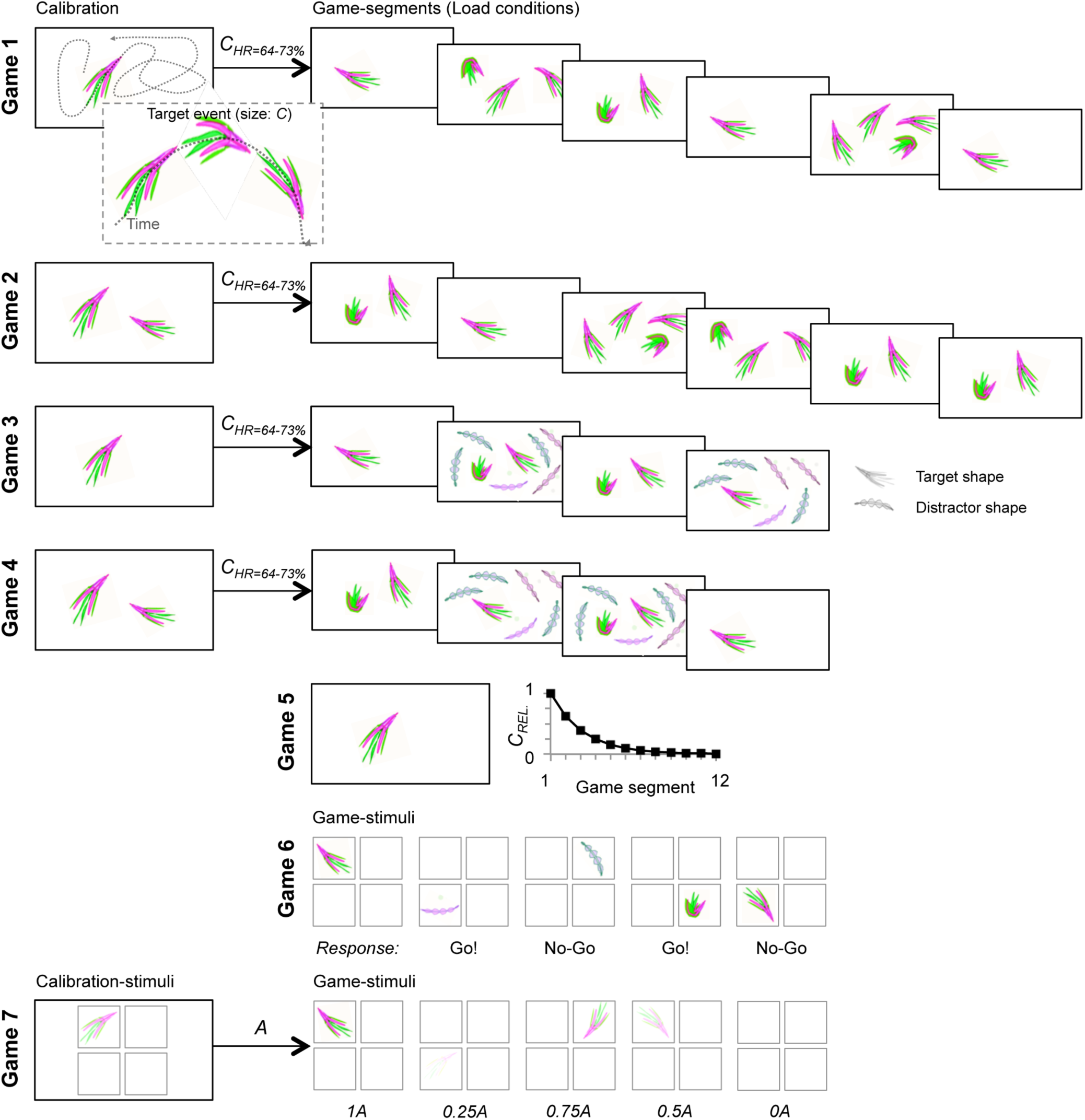
Schematic illustration of the training game task design and composition. For details, please see Materials and Methods section: *Training paradigm*. In short, the training program comprised seven different games, Games 1-7, that tapped primarily on visual acuity and contrast sensitivity in multiple attentional and working memory tasks. Subjects were presented with a pre-determined selection of games for each training day. The total training duration per week was ∼3.5 h, excluding the time spent on game parameter adaptation. In all tasks, the subject responded with a single keyboard-button press or withheld the response. **Games 1 and 2** were single- or multi-object visual tracking tasks where complex shaped objects moved along curved paths on screen and the subjects’ task was to respond whenever they observed a feature-change in any of the objects. Different game segments exhibited different numbers of to-be attended objects (attentional loads 1, 2, 3, and 4). Prior to each game, there was a calibration period with one (Game 1) or two (Game 2) objects during which the magnitude of the feature change (*C*) was adjusted to yield a detection rate (HR) of 64-73%. **Games 3 and 4** were visual-tracking games like Games 1 and 2 and had an identical calibration procedure and object mobility, but involved only attentional loads of 1 and 2, and exhibited in two out of four conditions six feature-wise distinct distractor objects to impose visual crowding. **Game 5** was a continuous single-object tracking task where the subjects reported the feature changes of a single object (as in Games 1-4). Game 5 had no calibration but rather started with very salient feature changes that in each of the 12 game segments decreased by a factor of 1.6 so that the subjects on average were able to reach segments 7-8 at a detection rate of >25%. **Game 6** was a Go/No-Go 1-back working memory task where the subject was presented stimuli with an object in one quadrant lasting ∼1 s at a rate of one stimulus in ∼2.5 s. The subjects task was to indicate whether the object in the current stimulus was different from the one in the previous stimulus regardless of quadrant and object rotation. **Game 7** was a threshold-stimulus-detection task where semi-transparent complex visual objects were presented randomly for 0.1 s and the subjects’ task was to report perceived stimuli. The object transparency was calibrated so that for an alpha-level A, detection rate of 0.5 was obtained at 0.5A. During the games, objects were at five equiprobable levels of A so that A were 0, 0.25, 0.5, 0.75, and 1.0.

### Visual acuity

Twenty-two of the 42 subjects (52.4%) showed a clinically relevant improvement in visual acuity [≥2 lines of vision (0.2 improvement in logMAR visual acuity)], and 7 of 42 subjects (16.6%) had improved to normal visual acuity (VA of logMAR 0 or better) during the study period (Table 3). Visual acuity significantly improved in the amblyopic eye in both treatment groups (Figure 3A). At the primary efficacy endpoint (10 weeks), the change in logMAR visual acuity from baseline was −0.167 [95% confidence interval (CI): −0.226 to −0.108] in the fluoxetine group and −0.194 (95% CI: −0.254 to −0.133) in the control group (both *p* < 0.001, Figure 3B). The marginal difference in visual acuity improvement between treatment groups was not statistically significant (*p* = 0.524). The visual acuity improvements observed in both groups were maintained 10 to 22 weeks after discontinuing all treatments (medication, eye patching, and perceptual training). In addition, visual gains of at least 0.2 logMAR units persisted in many subjects in both groups [9 of 20 fluoxetine subjects (45.0%), 6 of 14 control subjects (42.9%); Figure 3C].

**Table 3.** Summary of vision improvement in all 42 patients enrolled in the study, observed at any visit after the randomization visit.

| Summary of vision improvement<br>(at any visit after randomization visit) |  |  |  |
| --- | --- | --- | --- |
| Visual parameter | Test | Number of patients with improvement | Notes |
| Visual acuity | ETDRS | 22 / 42 (52%) | 7 / 42 patients (17%) achieved normal visual acuity ($<0.10$ logMAR). |
| Crowded near visual acuity | Landolt C ring | 9 / 42 (21%) | 9 patients improved to normal, defined as $\geq 0.7$ |
| Contrast sensitivity | Pelli-Robson | 2 / 2 (100%) | Two patients had abnormal contrast sensitivity at baseline. Both improved to normal contrast sensitivity, defined as $\geq 1.65$ log. |
| Binocularity | Bagolini striated glasses | 10 / 22 (45%) | Twenty-two patients had suppression or anomalous retinal correspondence at baseline. Ten patients changed to normal fusion. |
| Improve in at least one visual function test (during the whole course of the study) |  | 30 / 42 (71%) | Different patients improve on different visual parameters. No relation between visual acuity improvement and crowded near visual acuity, contrast sensitivity or binocularity. 6 / 42 patients improved in at least two visual parameters. |

**Figure 3.**
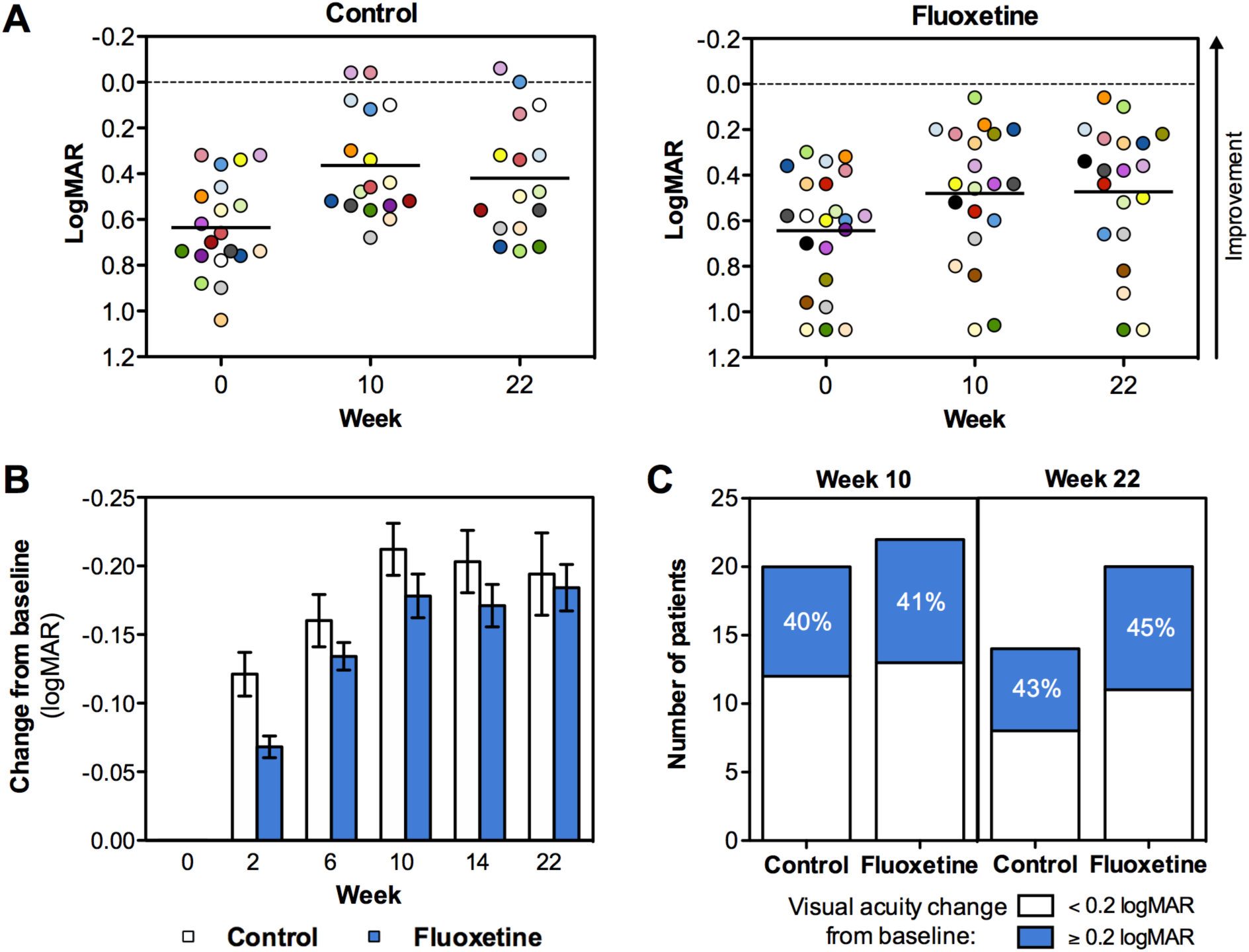
Improvement of visual acuity. (A) Scatter plots showing each individual patient’s visual acuity (amblyopic eye) at baseline, at week 10 (end of treatment/training) and at week 22 (end of follow-up), as measured by ETDRS chart (logMAR). Control group is shown on the left and fluoxetine group on the right. The limit of normal visual acuity (logMAR 0) is shown with a hatched line. (B) Average change in visual acuity from baseline as measured by ETDRS chart (logMAR) at baseline and after 2, 6, 10, 14 and 22 weeks. Average +/-95% CI in each timepoint when visual acuity was determined by ETDRS chart is shown. (C) Number of patients per group who showed improved visual acuity by ≥0.2 or <0.2 logMAR units at week 10 and 22 as compared to baseline.

### Binocularity, contrast sensitivity and crowded near visual acuity

Both treatment groups exhibited a positive response to treatment, with 30 of the 42 subjects (71.4%) showing an improvement in at least one visual function test (i.e., visual acuity, contrast sensitivity, binocularity, crowded near visual acuity) at any visit after the randomization visit. Binocularity, contrast sensitivity and crowded near visual acuity improved in both the control and fluoxetine groups. Binocular vision was assessed for both near (33 cm) and distant (4 m) vision. Twenty-two subjects (9 control group subjects, 13 fluoxetine group subjects) had suppression or ARC in the 4-m test at baseline (Figure 4A). At 10 weeks, the number of patients with suppression or ARC had decreased to 16 subjects (5 control group subjects, 11 fluoxetine group subjects). This change persisted through 22 weeks in 11 subjects (3 control group subjects, 8 fluoxetine group subjects). The results in the 33-cm test were similar for both groups (data not shown).

**Figure 4.**
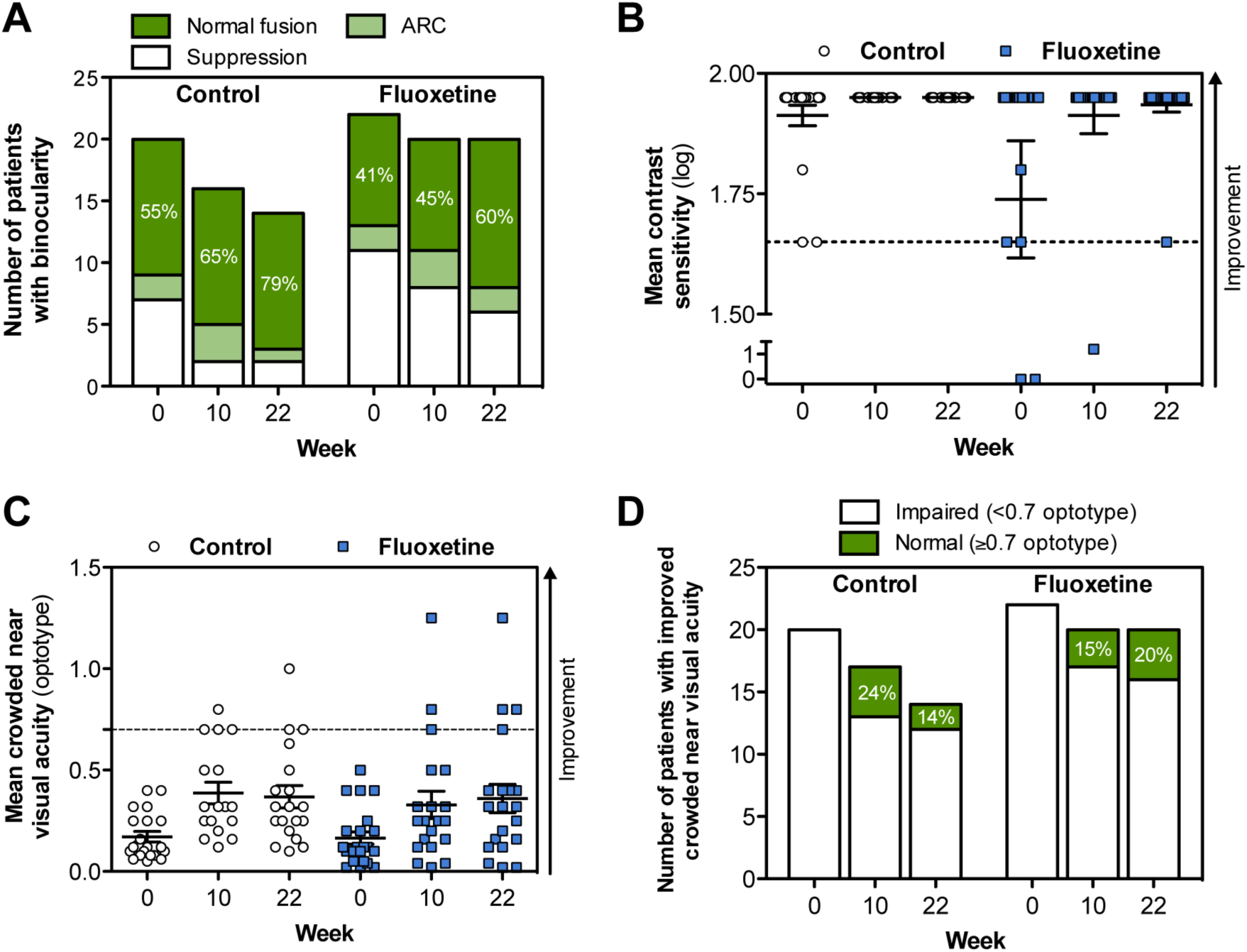
Change in binocularity, contrast sensitivity, crowded near visual acuity. (A) Number of patients per group who showed change in binocular vision (suppression, anomalous retinal correspondence (ARC) or normal fusion) at week 10 and 22 as compared to baseline. Bagolini striated glass test results at 4 meter distance are shown. (B) Mean contrast sensitivity (log value) as measured by Pelli-Robson chart. Normal contrast sensitivity (1.70) is indicated by a hatched line. Only two patients in the whole patient population (n=42) had significant contrast sensitivity impairment (i.e. 0 log) at baseline. Both patients received fluoxetine and improved to almost normal contrast sensitivity. (C) Crowded near visual acuity as measured by Landolt C ring charts. Normal crowded near visual acuity was considered as ≥0.7 (hatched line). In panels B and C, the direction of improvement is indicated by an arrow. (D) Number of patients per group who improved in crowded near visual acuity test at week 10 and 22 as compared to baseline.

> Contrast sensitivity was normal (1.70 log) at baseline in all but 2 fluoxetine subjects (4.8%) with severe amblyopia (both had 0 log values at baseline). Following fluoxetine treatment and perceptual training, both subjects had almost normal contrast sensitivity (10 weeks: 1.20 and 1.95 log, 22 weeks: 1.95 and 1.65 log; Figure 4B).

Crowded near visual acuity was assessed using Landolt C ring charts and was considered to be normal when 0.7 or smaller optotypes were detected. The baseline mean crowded near visual acuity in the amblyopic eye was 0.171 ± 0.116 in the control group and 0.165 ± 0.141 in the fluoxetine group.

Improvements in these values were observed after treatment in both study groups (Figure 4C). At 10 weeks, the control group had improved by 0.181 ± 0.027 (95% CI: 0.126 to 0.236) and the fluoxetine group had improved by 0.148 ± 0.026 (95% CI: 0.095 to 0.201, both p < 0.001). At 22 weeks, the control group had improved by 0.221 ± 0.036 (95% CI: 0.150 to 0.293) and the fluoxetine group had improved by 0.197 ± 0.033 (95% CI: 0.131 to 0.262, both p < 0.001). Figure 4D shows the distribution of patients with normal (≥ 0.7) and impaired (< 0.7) crowded near visual acuity.

The proportion of patients with an improvement in at least one of the visual function parameters was 60% in both treatment groups after 10 weeks of treatment. Three months after treatment completion (22 weeks), this proportion had increased to 70% in the fluoxetine group and 64% in the control group. Moreover, 73% of the fluoxetine group subjects and 70% of the control group subjects showed a clinically relevant improvement in at least one visual function parameter at some point during the study. Improvements in visual function parameters are summarized in Table 3.

Our clinical study protocol also included prespecified subgroup analyses, which assessed the effect of the severity of amblyopia (moderate/severe), binocularity (suppression/normal fusion), sex (female/male), compliance with software-based training (≥85%/<85%), and age (<40/≥40 years) on changes in visual acuity (logMAR). The subgroup analysis data are summarized in Table 4. Although the differences were not statistically highly significant (*p* = 0.048), there were more responders with normal fusion in the control group (6/11; 54.5%) than in the fluoxetine group (1/9; 11.1%), whereas there were more responders with suppression in the fluoxetine group (7/11; 63.6%) than in the control group (3/7; 42.9%). Finally, the visual acuity gains were associated with training compliance (*p* = 0.012) in both treatment groups (Table 4).

**Table 4.** Pre-specified subgroup analyses of visual acuity.

| Pre-specified subgroup analyses of visual acuity<br>(logMAR) |  |  |  |
| --- | --- | --- | --- |
| Subgroup variable | Interaction p-value<sup>1</sup> | Subgroup p-value<sup>2</sup> | Treatment p-value<sup>2</sup> |
| Severity of amblyopia<sup>3</sup> | p=0.774 | p=0.711 | p=0.633 |
| Binocularity (4 m)<sup>4</sup> | **p=0.048** | p=0.976 | p=0.544 |
| Gender<sup>5</sup> | p=0.161 | p=0.252 | p=0.493 |
| Training compliance<sup>6</sup> | p=0.873 | **p=0.012** | p=0.846 |
| Age<sup>7</sup> | p=0.678 | p=0.864 | p=0.525 |

The subgroup analysis results were obtained by incorporating the subgroup main effect and treatment-by-subgroup interaction effect into the primary RM ANCOVA model.
<sup>1</sup>Statistical significance of treatment-by-subgroup interaction in the FAS (LOCF) dataset
<sup>2</sup>Statistical significance of the subgroup and 10-week treatment effects in the FAS (LOCF) dataset from the model without the interaction effect (amblyopia severity, age, gender, compliance) or with the interaction effect (binocularity)
<sup>3</sup>Moderate/Severe
<sup>4</sup>Suppression/Normal fusion
<sup>5</sup>Female/Male
<sup>6</sup>≥85% / <85%
<sup>7</sup><40 / ≥40 years
Note: Severe amblyopia was more pronounced with younger, compliant, female and, especially, suppression patients.

### Safety

A total of 66 adverse events (AEs) were reported after initiating study treatments. Fifty-eight (87.9%) AEs occurred during treatment and 8 AEs occurred (12.1%) after treatment. Only 16 AEs (24.2%) were related or possibly related to the study treatment and all were reported during the treatment period. Eleven (16.7%) of these AEs occurred in the fluoxetine group and 5 (7.6%) occurred in the placebo group. None of the treatment-related AEs were reported following the 10-week treatment period and no AEs led to study withdrawal. One subject in the fluoxetine group exhibited transient mild diplopia that resolved spontaneously. Other reported AEs were not related to visual function. One serious adverse event (benign ovarian cyst of moderate severity) occurred during the study, but it was not related to study treatment.

## DISCUSSION

Amblyopia is a complex brain disorder that can restrict everyday life because of the visual limitations it imposes. Despite good screening programs and effective childhood treatments, amblyopia remains a common cause of lifelong visual impairment independent of location or ethnic origin.^4, 7, 9, 41^ The combination of adequate refractive correction and occlusion therapy (patching of non-amblyopic eye) has been the mainstay therapy for amblyopia of all etiologies. However, the benefits of various forms of occlusion therapy are greatest when therapy is started at an early age (<8 years). Therefore, early amblyopia detection and treatment is the most important factor for obtaining successful visual outcomes. Physiologically, the brain has the greatest plasticity during the critical period in early postnatal life. However, recent evidence strongly indicates that the primary sensory cortex may remain plastic into adulthood.^17–24, 27–30^ This finding suggests that there is a physiological basis for treating amblyopia in adulthood, which provides an opportunity to potentially alleviate this world-wide public health problem.

The current study examined whether fluoxetine, an SSRI known to modulate adult rat visual cortex plasticity^18^, can enhance the effects of patching/computer-based perceptual training combination therapy in adults with amblyopia. The treatment response was good in both the fluoxetine and placebo study groups, with an overall visual rehabilitation success rate of 52%. In addition, 17% of patients achieved normal visual acuity in the amblyopic eye (Table 3). Our results are in agreement with those of Li et al.^40^, who found that 33% of adult amblyopic subjects had a substantial improvement in visual acuity following video game-based perceptual training. In addition, our subjects had an average gain of approximately 2 lines of vision (0.2 logMAR), which was similar to improvements observed with other published training protocols.^38, 39^

The change in visual acuity after 10 weeks of study medication/perceptual training therapy was not significantly different between subjects taking fluoxetine and subjects taking a matching placebo. It may be that 20 mg of fluoxetine, the dose typically used to begin treatment of depression, was too small a dose to modulate neuroplasticity. It may also be that the training paradigm (new spectacles, patching, and computerized perceptual training) was so effective that the 20-mg dose of fluoxetine did not provide any additional benefit. Larger fluoxetine doses (up to 80 mg/day) are often used in depressed patients when the initial dose does not have the desired therapeutic effect. In addition, the 10-week treatment period may have been too short to maximize fluoxetine benefits and it is possible that a difference between treatment groups could have emerged after a longer treatment period.

Although there is evidence that some amblyopia treatments may be additive (optical correction in combination with patching or atropine)^32^, some reports have documented that all amblyopia treatment effects may not be additive. The effects of multiple amblyopia treatment paradigms were not synergistic in rodent models of amblyopia.^42^ Furthermore, environmental enrichment and fluoxetine treatment have been shown to induce similar levels of amblyopia recovery in rodents.^28^ Therefore, it is possible that perceptual training alone (with refractive correction) promotes the maximum amount visual cortex plasticity and that further treatments do not have additive benefits. The current study was not designed to determine this and future studies should include a group of subjects only treated with fluoxetine (no perceptual training). However, a rodent study found that fluoxetine alone had no effect on vision.^18^ Furthermore, there are no prior reports of visual benefits from fluoxetine monotherapy in amblyopic patients, even though millions of patients, and presumably thousands of amblyopic patients, have used the medication over the past three decades. In addition, a recent placebo-controlled, double-blinded, clinical study showed that 20 mg/day of fluoxetine for 19 days did not significantly affect visual perceptual learning in humans.^43^ These prior studies support the theory that 20 mg/day of fluoxetine may not be a large enough dose to effectively modulate visual cortex plasticity in adult humans.

Subjects in the current study received new spectacles with the proper refractive correction at baseline. Initiating the use of appropriate prescription glasses can improve amblyopia in children^44^ and adults^45^. Therefore, the use of new corrective lenses may have added to the visual gains observed in the current study. However, adults have lower plasticity than children so the effects of glasses may have been less prominent in our adult population. Furthermore, test-retest variability should be taken into account; this was low in the present study because visual acuity was measured under the same conditions and in the same locations by the same observers. Moreover, regression to the mean must be taken into account; because our study was placebo-controlled, the regression to the mean was reduced because both groups most likely exhibited an equal tendency.

We found that training compliance was well correlated with improved visual acuity. Therefore, the training software used in the current study could potentially be used in the clinical setting to personalize training and remotely monitor patient compliance. This would allow clinicians to adjust follow-up intervals based on treatment response rates. Understanding early treatment responses would also aid in determining which patients are likely to benefit from treatment and would allow for adjustment or discontinuation of treatment on the basis of individual responses. If no improvement is detected despite good compliance, treatment may be discontinued and the diagnosis of amblyopia may need to be reconsidered. Follow-up visit schedules can also be determined based on perceptual training performance.

Our study had several limitations. First, we did not have a true no-treatment control group. However, in both the placebo and fluoxetine groups, perceptual training compliance was well correlated with the magnitude of vision improvement. This finding strongly suggests that perceptual training resulted in neuroplasticity and subsequent visual benefits. Future clinical trials should include several control groups to examine the effects of individual interventions and their combined effects. Furthermore, the dose of fluoxetine and duration of its use should be varied in these studies. Second, the response to treatment was remarkably variable in both treatment groups. This may have resulted from the large amount of variation in amblyopia severity and etiology in our study population. Twenty-two subjects (52.4%) had abnormal binocularity at baseline, 10 of which had improvements in binocularity with study treatment. In addition, only 2 subjects with severe amblyopia had low contrast sensitivity at baseline. Contrast sensitivity deficits are sometimes found to correlate with the visual acuity in the amblyopic eye.^46–48^. Both subjects exhibited remarkable improvements in visual test results. Our results are in agreement with those of Zhou et al.,^49^ who showed that perceptual learning can improve visual acuity and contrast sensitivity in adult amblyopia patients. Nine of our subjects (21.4%) had improved crowded near visual acuity (Table 3). Hussain et al.^50^ found a significant association between reduction in crowding and visual acuity improvement in amblyopic adults. However, we did not find a significant correlation between visual acuity improvement and crowding, contrast sensitivity, or binocularity. It should be noted that the number of subjects in these subgroups was relatively small and that larger subgroups may have revealed additional statistically significant findings.

In conclusion, both fluoxetine and the software-based perceptual training were safe and well-tolerated, with fluoxetine treatment not offering further benefits over perceptual training. The training software used in the study simultaneously determined training compliance and improvements in visual function and showed that good training compliance is essential for treatment benefit. Therefore, software-based training, combined with eye patching, may improve visual function in adult patients with amblyopia. Furthermore, the computer program used here has the potential to become a robust tool for the treatment of amblyopia.

## MATERIALS AND METHODS

All study conduct adhered to the tenets of the Declaration of Helsinki and followed Good Clinical Practices. This study was reviewed and approved by the Regional Ethics Committee of Tampere University Hospital (centralized process for all centers in Finland) and the Research Ethics Committee of the University of Tartu (Estonia). Written informed consent was obtained from all subjects prior to performing any study examination or procedure. The study was registered in the European Clinical Trials Database (EudraCT) on October 1st, 2010 under the number 2010-023216-14.

This phase 2, multi-center, clinical study was a randomized, double-blind, placebo-controlled (for drug treatment), parallel-group trial performed to assess visual acuity improvement in the amblyopic eye, as measured by the Early Treatment of Diabetic Retinopathy Study (ETDRS) chart, following 10 weeks of medication (20 mg fluoxetine or placebo) and computer-based training (with the dominant eye patched). The following assumptions were made to calculate sample size: comparison of two equally sized groups, an intergroup difference (fluoxetine vs. placebo) in the change in logMAR visual acuity of at least 0.15, a standard deviation (SD) of 0.15, and a subject drop-out rate of 10%. Thirty-four subjects needed to be randomized to power the study to 80%, assuming a two-sided type I error rate of 5%.

### Study subjects

Four eye clinics in Finland and Estonia enrolled 42 subjects between June 2011 and April 2013. Study inclusion and exclusion criteria are fully described in Table 1. Briefly, adult patients with monocular amblyopia with no other ocular or neurological abnormalities were considered for enrollment. Included subjects were 19 to 57 years of age and had moderate (0.3–0.6 logMAR difference) to severe (>0.6 logMAR difference) amblyopia due to myopic or hyperopic anisometropia (≤4.25 D) or congenital esotropia. The lower limit for anisometropia was not set in the study protocol. The investigators considered the amblyopia to be of the anisometropic type if no strabismus had been diagnosed in childhood and the refractive error was at least 1 D of anisometropia, determined as the spherical equivalent, in childhood. Patients with other primary forms of strabismus, extrafoveal (eccentric) fixation, or who used antidepressant drugs in the past 6 months were excluded.

### Study examinations

Eligibility, demographic data, medical history, relevant medication, vital signs, physical examination, blood and urine samples (including urine pregnancy test for fertile women), and amblyopia were assessed at screening. Amblyopia was confirmed at screening and was defined as an interocular ETDRS best-corrected visual acuity difference of at least two lines and/or a logMAR visual acuity between 0.30 and 1.10 in the amblyopic eye and 0.10 or better in the dominant eye. Prior to randomization, patients received new spectacles based on non-cycloplegic refraction to ensure best-corrected vision during the study.

A thorough ophthalmic examination was conducted at each of the seven scheduled visits (at weeks −2 (screening), 0 (randomization), 2, 6, 10, 14 and 22; Figure 1) during the 26-week study period. Vision tests included the assessment of binocularity, visual acuity, crowded near visual acuity, and contrast sensitivity and were performed with the refractive error corrected. In addition, presbyopic correction was used for crowded near visual acuity testing in presbyopic subjects.

Binocularity was examined using the Bagolini striated glass test^51^ before monocular testing. Lens striations were placed at 135° before the right eye and 45° before the left eye using lorgnette frames. This testing setup allows each eye to receive the same fusible image with each fixation streak oriented perpendicular to the striations and 90° away from the other eye. The test enables the evaluation of simultaneously perceived images with a minimal dissociative effect and it was performed at near (33 cm) and distance (4 m) under normal lighting conditions. Binocularity was categorized as suppression (1 light and only 1 line were seen), normal fusion (binocular single vision, BSV; 2 lines were seen as X and 1 light at the center), anomalous retinal correspondence (ARC; harmonious if 1 light and 2 lines were seen, but one of the lines was broken due to foveal suppression, or inharmonious if 1 light and 2 lines were seen, but the lines did not cross at the center where the light was located) or diplopia (2 lights and 2 lines were seen). However, none of the subjects had diplopia in the current study.

Visual acuity was assessed under standardized lighting conditions (self-calibrated test lighting with a constant light level of 85 cd/m^2^) using a large-format standardized ETDRS light box (ESV3000 with LED lights, VectorVision, Greenville, OH) placed 4 meters from the subject. Three different ETDRS charts (charts R, 1 and 2) were used to prevent subjects from memorizing eye charts. Visual acuity was assessed in the amblyopic eye first and was measured as the number of correctly identified letters. A clinically relevant visual acuity improvement was defined as a 0.2 or greater decrease in the logarithm of the minimum angle of resolution (logMAR) visual acuity (i.e., 2 lines or 10 characters on the ETDRS chart).

Contrast sensitivity was determined under standardized lighting conditions using a Pelli-Robson chart at a distance of 1 m (charts A and B), using previously established age-dependent normative values.^52^ Crowded near visual acuity was assessed using a specific crowded Landolt C ring chart booklet at a distance of 40 cm.^53^ Crowded near visual acuity was defined by the smallest line in which the subject correctly identified at least 8 of 12 letters (≥66.7%). The right eye was tested first in all tests requiring charts. The contralateral eye was occluded during testing and charts were switched between eyes. All eye examination test charts and Bagolini striated glasses tests were standardized and validated for trial endpoint measurement. All staff that evaluated vision were masked to subject group assignment.

Treatment safety was assessed using ophthalmoscopy, biomicroscopy, intraocular pressure (IOP) measurement, laboratory safety tests [hematology (hemoglobin, hematocrit, erythrocyte count, leukocyte count, platelet count), clinical chemistry (alanine aminotransferase, alkaline phosphatase, aspartate aminotransferase, creatinine, gamma-glutamyl transferase, potassium, sodium, urea), and urine analysis (blood, glucose, ketones, protein, pH)], vital signs, and physical examination performed at screening and at each study visit. Adverse events and changes in concomitant medications were recorded at each study visit.

### Study medication

Fluoxetine capsules were manufactured by Orion Corporation (Espoo, Finland) and the matching placebo capsules were manufactured by Corden Pharma GmbH (Plankstadt, Germany). Study subjects were randomly assigned to receive either 20 mg fluoxetine each day (hard capsule) or a matching placebo. Randomization was done in a 1:1 fashion in blocks of 4 and was stratified by site. Randomization was also stratified by amblyopia severity, determined using interocular visual acuity difference (moderate: 0.3-0.6 logMAR difference, severe: >0.6 logMAR difference). The randomization structure was designed by a biostatistician and the final randomization list was generated by an independent person who had no contact with study subjects or study data. Medication was pre-packed and serially numbered so that subjects were assigned to a study group by giving them the next available medication number in the sequence. A drug accountability log was maintained by study-authorized personnel. The receipt, dispense and return of study medication was recorded in this log. Patients were instructed to return dispensed medication bottles at the next visit, even if the bottles were empty. The number of capsules dispensed and returned was reconciled against the number of days between the visits and any discrepancies were accounted for. After 10 weeks of receiving study medication, subjects were weaned off the daily medication (1 capsule every other day for the next 2 weeks, Figure 1B).

### Perceptual training

All subjects completed daily computerized training with eye patching during the 10-week period of receiving study medication (Figure 1A). The principle underlying the perceptual training software developed for this study is fully described in the Electronic Supplementary Materials and is illustrated in Figure 2 and Supplementary video 1. All subjects received new spectacles before randomization and were instructed to wear an eye patch over their dominant eye while performing daily computerized perceptual training. The training software was used to track training compliance, which was calculated by dividing the total accomplished training time with the total prescribed training time. Training compliance was automatically reported to the study site prior to each scheduled visit.

All subjects were given an eye patch and were instructed to wear it over the non-amblyopic eye for 1 hour each day. Subjects were also instructed to complete approximately 30 minutes of the computer-based training each day while they were wearing the patch and their spectacles.

The training period was divided into ten 1-week segments and each subject played an identical composition of games each week. The maximum total training time over the 10-week training period was 35 hours. The training program was made up of seven different games wherein the performance was primarily determined by visual acuity and contrast sensitivity and secondarily by attention and mental effort. Thus, the training was primarily focused on visual acuity and contrast sensitivity and was aimed at their improvement. A schematic illustration of the training game task design and composition is shown in Figure 2. In addition, the computerized training setup, training protocol structure, and individual training game design are described in detail in the Electronic Supplementary Materials. Data on behavioral performance were collected on a per-game basis. For each game type, the corresponding weekly test outcome measures were obtained by pooling the data from all individual games of that type played in that week (see Electronic Supplementary Materials).

### Study outcomes

The primary outcome of the study was an improvement in visual acuity in the amblyopic eye, as measured by the ETDRS chart, from baseline (week 0, randomization) to the 10-week visit (end of treatment). Secondary outcomes included the change from baseline in binocularity, contrast sensitivity and crowded near visual acuity to week 10 week. The persistence of changes observed at 10 weeks was also evaluated at the end of the follow-up period (week 22). Treatment safety was assessed using adverse event incidence and ophthalmological examination findings throughout the study. Exploratory outcomes included changes from baseline in training measures at each study visit.

### Data analyses

The primary analysis population was the full analysis dataset (FAS), which included all randomized patients who had received at least one dose of study medication (intention-to-treat principle). A last observation carried forward (LOCF) imputation was applied up to week 10 for subjects who did not complete the study and those who were non-compliant with the treatment.

Differences in visual acuity and crowded near visual acuity between the two treatment groups were evaluated using the repeated measurements of analysis of covariance (RM ANCOVA) method with baseline values as a covariate. The model included the study center, treatment and time point (visit) as main effects, and treatment by time point (visit) as an interaction effect. With regard to the primary endpoint, differences between the treatment groups with regards to change in the logMAR visual acuity at 10 weeks (and a 95% CI for the difference) were estimated using RM ANCOVA models with a contrast. A secondary RM ANCOVA analysis was used to compare the least square means between the treatment groups at the end of the follow-up (22 weeks) period to determine if treatment effects were maintained. Contrast sensitivity was not analyzed with RM models because of low variability in the dataset. Pearson’s chi-square test was used to compare categorical variables. Some binocularity categories contained a low number of subjects. Therefore, differences between treatment groups in binocularity were evaluated using Fisher’s exact test.

## ACKNOWLEDGEMENTS

The authors thank prof. Lamberto Maffei for his input to the design of the study.

## ADDITIONAL INFORMATION

### Author Contributions

HJH, MP, LL, SP, SB, EC and HU designed research; HJH, MP, LL, SP, VS, EK, JL, MLL, SB performed research and analyzed the data; HJH, MP, JL, EC and HU wrote the paper.

